# LINE-1 transposon derepression and epigenetic remodeling in Retinoblastoma

**DOI:** 10.64898/2026.07.25.740688

**Authors:** Aman Verma, Sugam Kumar Patel, Akhil Varshney, Pratyashaa Paul, Ria Sachdeva, Virender Singh Sangwan, Sima Das, Bhavana Tiwari, Anil Tiwari

**Affiliations:** Eicher Shroff-Centre for Stem Cell Research, Dr. Shroff’s Charity Eye Hospital, Delhi; Oculoplasty and Ocular Oncology Services, Dr. Shroff’s Charity Eye Hospital, Delhi; Centre for Doctoral Studies, Manipal Academy of Higher Education, Manipal; Department of Ocular Genetics, Centre for Unknown and Rare Eye Diseases, Dr Shroff’s Charity Eye Hospital, Daryaganj, New Delhi, India; Department of Ophthalmology, University of Loma linda, USA; Department of Biological Sciences, Indian Institute of Science Education and Research Berhampur, India

**Keywords:** RB1 tumor suppressor, L1/LINE-1, Pediatric cancer, Transposable elements, Retinoblastoma

## Abstract

Retinoblastoma (RB), the most common pediatric intraocular malignancy, is initiated by biallelic inactivation of the RB1 tumor suppressor gene. Although LINE-1 (L1) retrotransposon activation has been linked to genomic instability and tumor evolution in several adult cancers, its contribution to pediatric malignancies remains poorly understood. Here, we investigated L1 expression in retinoblastoma samples from an Indian pediatric cohort by integrating transcriptomic, locus-specific, and immunohistochemistry. Transcriptomic profiling revealed increased L1 expression in RB tumors compared with control retinal tissues, which was validated by ORF1p immunostaining in tumor sections. Locus-specific analysis identified a subset of transcriptionally active L1 loci, suggesting selective activation of young L1 elements. We also observed altered expression of genes associated with proliferation, chromatin regulation, and oncogenic signaling, including MYCN, MDM2, E2F3, SOX4 and CREBBP. Our findings identify previously underexplored L1 activation as a molecular feature associated with retinoblastoma and suggest that disruption of transposon silencing accompanies the RB transcriptional landscape. Validation in larger patient cohorts will be important; however, this study provides a foundation for exploring L1 activity as a potential molecular marker and future therapeutic vulnerability in retinoblastoma.

**Highlights:**

- Transcriptomic profiling reveals increased LINE-1 expression in retinoblastoma.
- ORF1p protein is robustly expressed in retinoblastoma.
- A subset of LINE-1 loci exhibits transcriptional activation, suggesting locus-specific derepression.

**In Brief:** Retinoblastoma is initiated by RB1 loss, but the contribution of transposable elements to its molecular landscape remains poorly understood. Here, we show that retinoblastoma is associated with LINE-1 promoter hypomethylation, increased LINE-1 transcript expression, ORF1p accumulation, and preferential activation of selected LINE-1 loci. These changes occur together with altered expression of oncogenic signalling pathways and host factors involved in transposable element repression, identifying dysregulated LINE-1 control as a molecular feature of this paediatric cancer.

**Graphical Abstract:** 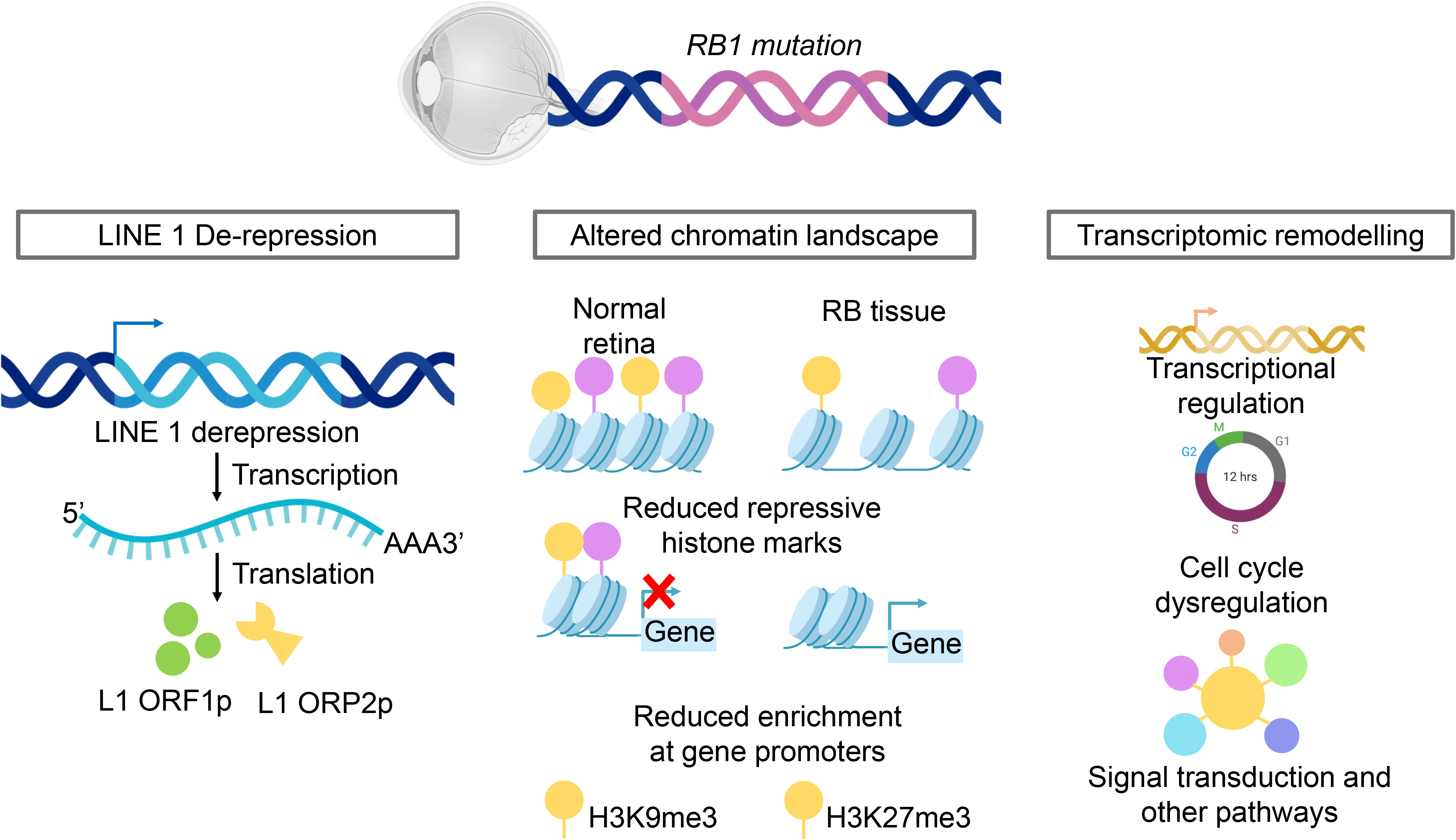

Retinoblastoma exhibits epigenetic activation of LINE-1 characterized by promoter hypomethylation, increased expression of young LINE-1 elements, and ORF1p accumulation. This is accompanied by coordinated transcriptomic remodeling and disruption of host mechanisms that normally restrain transposable elements, linking LINE-1 activation to the molecular landscape of retinoblastoma.

## Introduction

Retinoblastoma (RB) is a rare but aggressive form of paediatric eye cancer, primarily caused by biallelic inactivation of the *RB1* gene (1,2). Most bilateral tumors are caused due to germline mutations of the *RB1* gene (3). Although early diagnosis and treatment lead to high survival rates in high-income countries, significant disparities persist in low-and middle-income countries such as India, where patients often present with advanced disease (1,4,5). Despite genetic similarities to RB cases reported worldwide, Indian RB patients remain underrepresented in molecular studies, limiting our understanding of region specific disease biology (6,7).

The RB1 gene is best known for its pivotal role in regulating the G1/S transition of the cell cycle (8), but it also functions as an important epigenetic regulator. Beyond cell-cycle control, RB1 interacts with chromatin modifying complexes to maintain transcriptional repression of repetitive genomic elements, including transposable elements (TEs), thereby contributing to genome stability (9). Loss of RB1 function interrupts with these repressive mechanisms, resulting in widespread epigenetic alterations and increased chromosomal instability (10). Aberrant activation of transposable elements has emerged as a common feature of many human cancers, where these genetic elements represent an endogenous source of genomic instability(11) .Among these, Long Interspersed Nuclear Element-1 (LINE-1 or L1) is the only autonomously active retrotransposon in the human genome (12). LINE-1 propagates through a copy-and-paste mechanism in which its RNA transcript is reverse transcribed and inserted into new genomic loci (13,14). In addition to generating new insertions, LINE-1 activity can promote DNA damage, chromosomal rearrangements, transcriptional dysregulation, and activation of innate immune pathways, collectively contributing to genome instability and tumor evolution (15–19). RB1 mutations that results into loss of function alters heterochromatin landscape and epigenetic silencing, suggesting that RB1 deficiency may promote the derepression of transposable elements, including LINE-1 (20). This plausible connection is also supported by studies demonstrating that the RB1 pathway contributes to the transcriptional repression of repetitive genomic elements (21) .RB1 cooperates with chromatin modifiers such as HDACs, SUV39H1, and EZH2 to promote the deposition and maintenance of repressive chromatin marks, including H3K9me3 and H3K27me3, at repetitive sequences (22,23). Loss of RB1 has been associated with increased heterochromatin decompaction (24), altered histone modifications, and increased expression of repetitive elements in multiple experimental systems (25). Although direct casual mechanisms between RB1 mutation to LINE-1 activation in retinoblastoma remains limited, these observations provide a rationale for investigating whether disruption of RB1-dependent epigenetic changes contributes to LINE-1 derepression and its role in RB.

Despite this framework, to the best of our knowledge, no previous study has directly examined L1 expression in RB tumors from Indian patients. Therefore, our study fill this gap and demonstrate L1 expression in Indian RB tumors using transcriptomic studies and validation using immunohistochemistry. Our results show that upregulation of L1 transposons at both the RNA and protein levels. We observed increased expression of genes associated with proliferation, chromatin regulation, and oncogenic signaling, including MYCN, MDM2, E2F3, SOX4 and CREBBP. We demonstrate patient specific expression patterns of LINE-1 loci, with a subset of loci selectively expressed in retinoblastoma tumors. These findings suggest reactivation of LINE-1 elements that are otherwise expected to remain transcriptionally silent. Our cohort size is limited, however, to our knowledge these results demonstarte the first transcriptomic evidence of LINE-1 dysregulation in retinoblastoma from an Indian patient cohort.

This study underscores the significance of understanding to transposon biology in pediatric cancers and emphasises the potential of L1 as a molecular marker or therapeutic target in RB. Future studies with larger cohorts and integrated epigenomic data will be required to fully dissect out the mechanisms driving TE activation and its role in RB progression.

## Results

### Result 1: Clinicopathological characterization of the retinoblastoma cohort

To understand the molecular and epigenetic landscape of retinoblastoma (RB) in Indian patients, we assembled a cohort of 30 enucleated RB tumors that were collected from patients undergoing surgical treatment. These samples were used across multiple experiments, where bisulphite sequencing was performed to examine LINE-1 promoter methylation analysis (n=9), RNA sequencing (RNA-seq; n=6), quantitative Real time PCR (qPCR; n=17) for quantifying L1 transcripts, and finally validation using immunohistochemistry (IHC; n=13) where ORF1p, a L1 transposon encoded protein was stained using ORF1p antibody, with several samples, here n indicate sample number, analyzed by more than one methodology to ensure cross-validation of our observations (**Figure 1A**).

**Figure 1.**
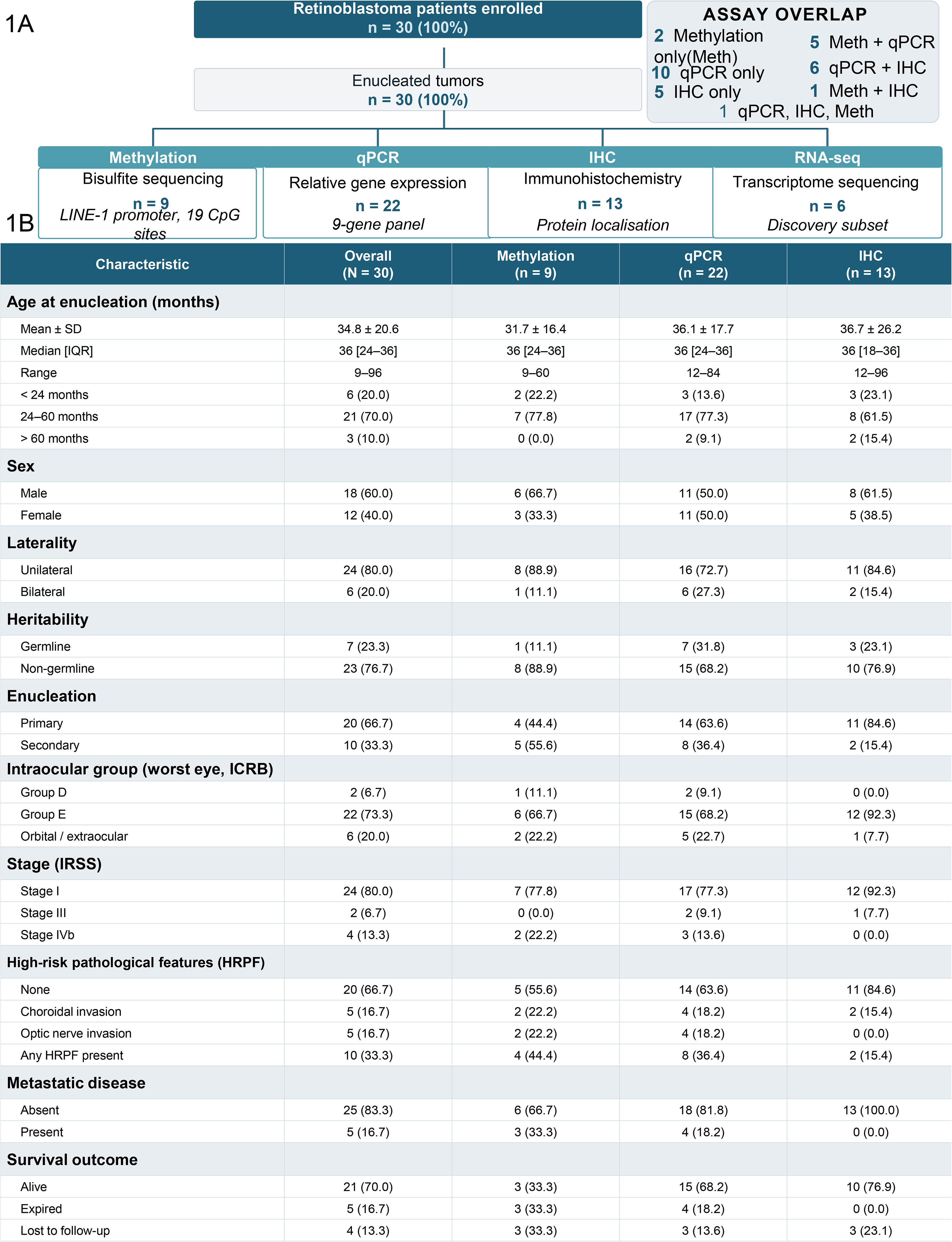
Clinicopathological characterization of the retinoblastoma cohort and analysis of LINE-1 promoter methylation. **A** Overview of the study cohort showing patient recruitment (N = 30) and allocation of samples for LINE-1 promoter methylation analysis (n = 9), quantitative RT-PCR (n = 21; LINE-1 measurements available in n = 17), immunohistochemistry (n = 13), and RNA sequencing (n = 6). Assay groups are not mutually exclusive. **B.** Demographic characteristics of the retinoblastoma cohort and assay subgroups, including age at enucleation, sex, laterality, germline status, and primary versus secondary enucleation. Data are presented as n (%) or median [IQR], as appropriate.

The clinicopathological characteristics of the patient cohort are summarized and presented in **Figure 1B**. The mean age at diagnosis was 34.8 ± 20.6 months (median, 36 months), with most patients presenting between 24 and 60 months of age. The cohort comprised 18 males (60%) and 12 females (40%), and unilateral disease was observed in 80% of cases. Most tumors were classified as non-heritable (76.7%), whereas 23.3% of patients carried germline RB1 mutations. Primary enucleation was performed in 66.7% of patients, while 33.3% underwent secondary enucleation following prior treatment.

Consistent with the advanced disease stage at presentation, the majority of tumors were classified as International Classification of Retinoblastoma (ICRB) Group E (73.3%), and 80% of patients were diagnosed with International Retinoblastoma Staging System (IRSS) Stage I disease. High-risk pathological features, including choroidal and/or optic nerve invasion, were identified in one-third of tumors, whereas metastatic disease was observed in 16.7% of patients. At the time of last follow-up, 70% of patients were alive, 16.7% had died, and 13.3% were lost to follow-up. We selected clinically well characterized specimens from the retinoblastoma cohort for molecular analysis. LINE-1 promoter methylation was assessed by bisulfite methylation analysis, while LINE-1 and transposable element expression were examined using transcriptomic profiling and RT-PCR based validation. In addition, immunohistochemistry was performed to validate L1-ORF1p protein expression in tumor tissues.

### Result 2: Retinoblastoma exhibits LINE-1 promoter hypomethylation

Among the host mechanisms that maintain transposable element repression (26), DNA methylation of the LINE-1 (L1) 5′ untranslated region (5′UTR) is a major mechanism for transcriptional silencing of endogenous LINE-1 elements in somatic tissues. To investigate whether this epigenetic repression is altered in retinoblastoma, we assessed LINE-1 promoter methylation in primary retinoblastoma tumors (n=6) and control retinal tissues obtained from patients undergoing enucleation or evisceration for non-retinoblastoma ocular conditions (n = 3). A schematic representation of the full-length human LINE-1 element, including the 5′UTR, ORF1, ORF2, 3′UTR, and the nineteen CpG dinucleotides analyzed in this study, is shown in **Figure 2A**.

**Figure 2:**
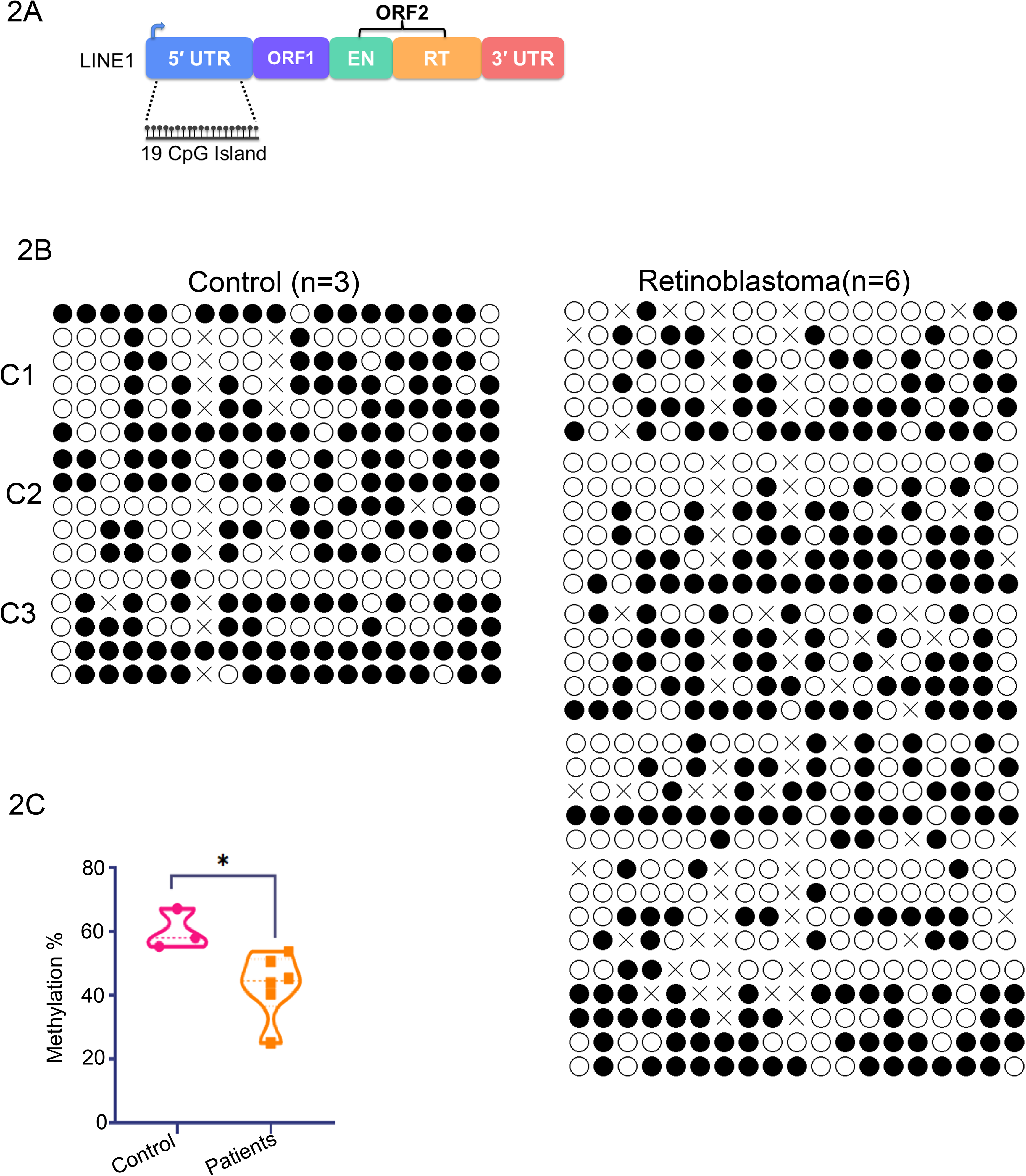
LINE-1 promoter hypomethylation in retinoblastoma. **A.** Schematic representation of a full-length human LINE-1 (L1Hs) element showing the 5′ untranslated region (5′UTR), ORF1, ORF2, and the 3′UTR. The positions of the nineteen CpG sites analyzed within the LINE-1 promoter are indicated. **B.** Bisulfite sequencing analysis of LINE-1 promoter methylation in control retinal tissues (n = 3) and primary retinoblastoma tumors (n = 6). Approximately four independent clones were analyzed per sample. Each row represents an individual clone. Filled circles indicate methylated CpG sites, open circles indicate unmethylated CpG sites, and crosses denote mutated CpG sites. CpG positions are shown relative to the human L1Hs consensus sequence. **C.** Quantification of LINE-1 promoter methylation in control retinal tissues (n = 3) and retinoblastoma tumors (n = 6). Data represent the percentage of methylated CpG sites per sample and are presented as mean ± SEM. Statistical significance was determined using a two-tailed Mann–Whitney U test..

We found methylation of the LINE-1 promoter in control retinal tissues using bisulfite sequencing, consistent with transcriptional silencing of endogenous LINE-1 elements in the normal retina. In contrast, LINE-1 promoter hypomethylation was observed across the nineteen CpG sites in retinoblastoma patient samples (**Figure 2B**). The extent of methylation varied among individual retinoblastoma tumor samples, but reduced LINE-1 promoter methylation was observed across the retinoblastoma cohort.

Quantification of methylation across all CpG sites and samples showed a reduction in overall LINE-1 promoter methylation in retinoblastoma compared with control retinal tissues (**Figure 2C**). These results suggest that the LINE-1 promoter undergoes epigenetic derepression in retinoblastoma, providing a possible molecular basis for aberrant LINE-1 activation. Since LINE-1 promoter hypomethylation is a well-established prerequisite for its transcriptional activation, we next examined whether reduced promoter methylation was associated with increased LINE-1 expression in retinoblastoma.

### Result 3. LINE-1 promoter hypomethylation is accompanied by transcriptional activation of evolutionarily young LINE-1 elements in retinoblastoma

To examine if LINE-1 promoter hypomethylation is linked to transcriptional activation of endogenous retrotransposons, we performed RNA sequencing of primary retinoblastoma tumors (n=6) and control retinal tissues (n=3). Principal component analysis (PCA) showed a clear separation between retinoblastoma and control samples, with principal component 1 (PC1) accounting for 68% of the total variance and principal component 2 (PC2) explaining an additional 10%, indicating distinct global transcriptomic profiles between the two groups (**Figure S1A**). Analysis of transposable element (TE) families revealed widespread dysregulation extending beyond LINE-1, including LINEs, SINEs, LTR retrotransposons, and other repetitive elements (**Figure 3A**).

**Figure 3:**
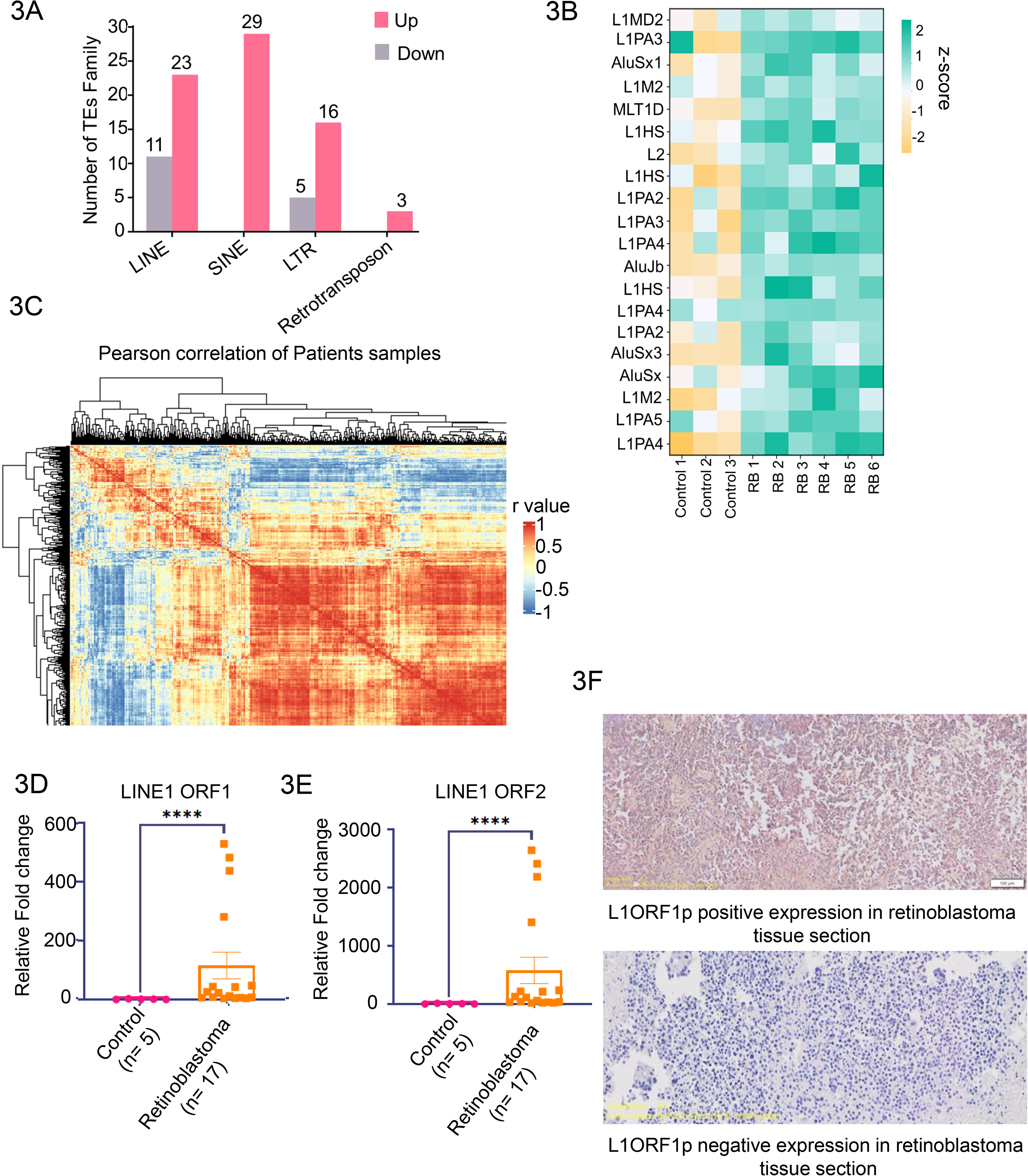
LINE-1 transcriptional activation and ORF1p expression in retinoblastoma. **A.** The bar plot represent total number of up and down regulated TEs family of LINEs, SINEs, LTRs, and retrotransposons . The x-axis represents the different TEs and y-axis represents TE family. **B.** Heatmap showing the twenty most highly expressed LINE-1 (L1) and Alu elements in control retinal tissues and primary retinoblastoma tumors. Each row represents an individual transposable element, and each column represents an individual sample. Relative expression is displayed as a normalized Z-score. The colour codes ranges from green to yellow, green colour shows up regulated and yellow represents downregulated and intermediate shades shows no change. **C.** Pearson correlation analysis of differentially expressed young LINE-1 subfamilies (L1HS, L1PA2, L1PA3, L1PA4, and L1PA5) and Alu families across six retinoblastoma RNA-seq samples. Hierarchical clustering identified positively and negatively correlated transposable element clusters, shown in red and light blue, respectively. **D and E.** Quantitative RT-PCR analysis of LINE-1 ORF1 (D) and ORF2 (E) transcript expression in control retinal tissues (n = 5) and primary retinoblastoma tumors (n = 17). Transcript levels were normalized to GAPDH and expressed relative to controls. Data are presented as mean ± SD. ****p<0.0001, statistical significance was determined using a two-tailed Mann-Whitney U test. **F.** Representative immunohistochemical staining of LINE-1 ORF1p in primary retinoblastoma tissues using an anti-ORF1p antibody. The upper panel shows positive ORF1p staining in tumor tissue. The lower panel shows negative staining in adjacent tissue serving as an internal negative control. ORF1p positive cells are visualized by brown DAB chromogenic staining, and nuclei are counterstained with hematoxylin (blue). Whole-slide images were acquired using an Olympus OlyVIA scanner at 20× magnification. 100µm.

We observed a greater number of TEs exhibiting increased rather than decreased expression in retinoblastoma relative to control retina. Among the dysregulated elements, evolutionarily young LINE-1 subfamilies were prominently represented. Heatmap analysis of the twenty most highly expressed LINE-1 and Alu elements clearly separated retinoblastoma from control samples and demonstrated increased expression of multiple L1HS, L1PA, and Alu elements in tumors (**Figure 3B and S1B**). Pearson correlation analysis further revealed coordinated expression patterns among young LINE-1 subfamilies (L1HS, L1PA2, L1PA3, L1PA4, and L1PA5), identifying distinct positively and negatively correlated TE modules across retinoblastoma samples (**Figure 3C**).

To validate these transcriptomic findings, we quantified LINE-1 expression in an independent cohort of control retinal tissues (n=5) and primary retinoblastoma tumors (n= 17). Quantitative RT-PCR demonstrated significant upregulation of both LINE-1 ORF1 and ORF2 transcripts in retinoblastoma compared with control retina (**Figure 3D and E**). Consistent with increased transcript abundance, immunohistochemistry staining experiments revealed robust ORF1p expression in 12 of 13 retinoblastoma specimens, whereas only a single tumor lacked detectable staining, a representative image is shown in (**Figure 3F**), confirming that LINE-1 transcriptional activation is accompanied by accumulation of LINE-1 protein.

Because only a subset of genomic LINE-1 copies retain transcriptional and retrotransposition potential, we next investigated LINE-1 expression at single-locus resolution. The L1 analysis at locus level demonstrated that expressed LINE-1 loci belonged predominantly to evolutionarily young subfamilies, with L1PA2, L1PA3, and L1PA4 accounting for the largest proportion of detected loci, followed by L1HS (**Figure 4A**). Heatmap analysis identified 42 differentially expressed L1HS loci that clearly distinguished retinoblastoma from control samples (**Figure 4B**). Similarly, analysis of 548 loci belonging to the L1PA2, L1PA3, and L1PA4 subfamilies revealed widespread heterogeneity in locus-specific expression, with numerous loci consistently upregulated across retinoblastoma tumors (**Figure 4C**). These findings indicate that LINE-1 activation in retinoblastoma is not restricted to a small number of loci but involves coordinated but selective activation of multiple evolutionarily young LINE-1 copies distributed throughout the genome. These observations suggests that LINE-1 promoter hypomethylation in retinoblastoma is accompanied by robust transcriptional activation of evolutionarily young LINE-1 elements at both the family and individual locus levels, resulting in increased LINE-1 RNA and ORF1p protein expression.

**Figure 4.**
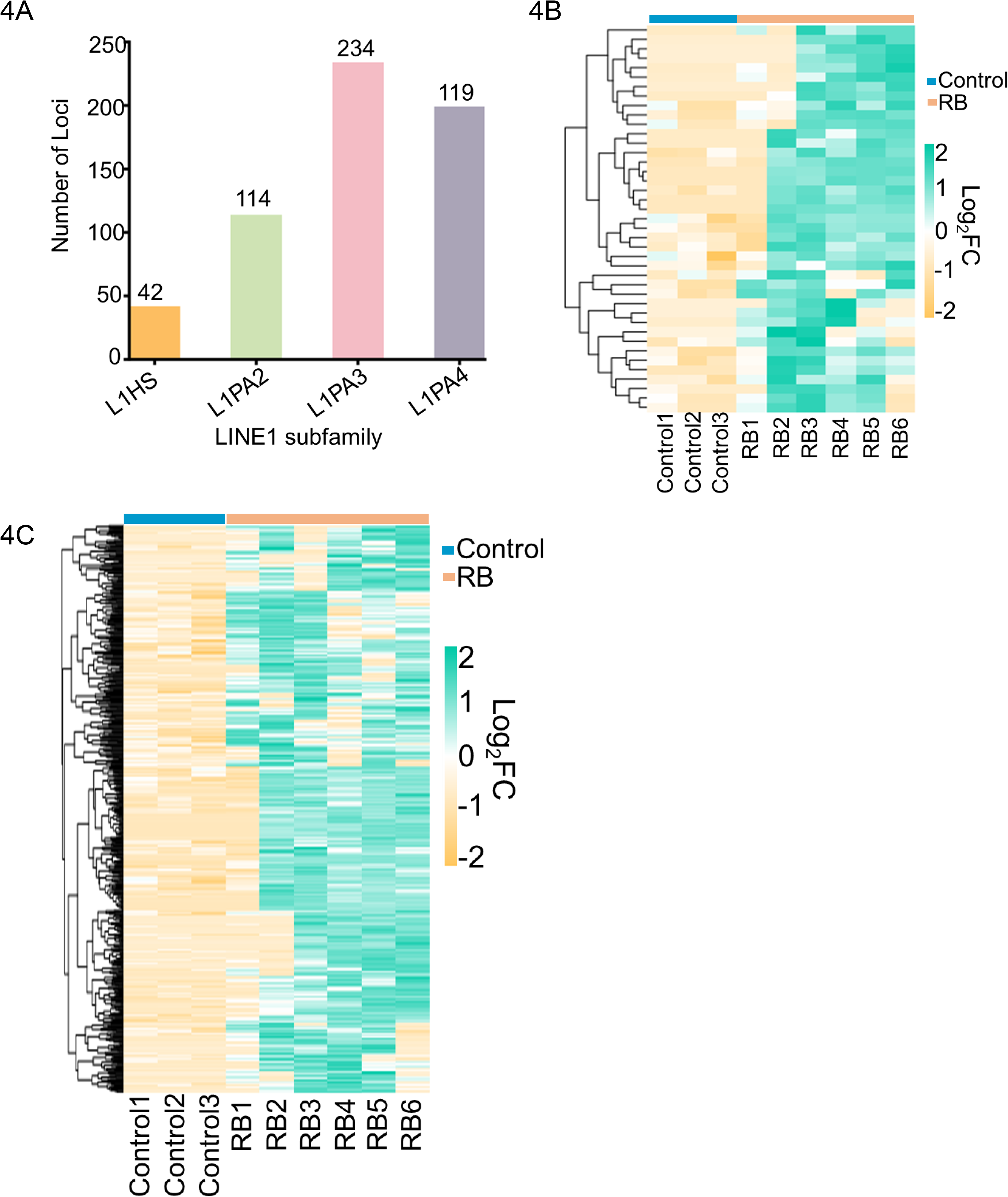
Evolutionarily young LINE-1 loci are activated in retinoblastoma. **A.** The bar plot represents total number of LINE1 subfamily loci present in the control and RB patient’s sample. The x-axis shows the LINE 1 subfamily level and y-axis represent the total number of LINE1 Loci. **B.** The heatmap represents total 42 L1HS loci of control vs RB patient’s sample. The x-axis represents shows the sample group and the y-axis individuals L1HS loci,. The colour codes ranges from green to yellow. The green colour represents up regulated, yellow colour shows down regulated and white color shows no change. Relative expression is displayed in log_2_FC. **C.** The heatmap represent the 548 loci of LIPA2, LIPA3, LIPA4 in control vs RB patient’s sample. The x-axis represents sample group, and the y-axis shows the LINE1 loci. The colour codes ranges from green to yellow. The green colour represents up regulated, yellow represents down regulated and white colour shows no change. Relative expression is displayed in log_2_FC.

### Result 4. LINE-1 activation is accompanied by widespread transcriptional changes of oncogenic signalling pathway

To investigate the transcriptional consequences associated with LINE-1 activation in retinoblastoma, we analysed RNA sequencing of primary retinoblastoma tumours (n=6) and control retinal tissues (n=3). Differential expression analysis identified extensive transcriptomic remodelling, with 3,622 genes upregulated and 1,994 genes downregulated in retinoblastoma, indicating a global shift in gene expression associated with tumour development (**Figure 5A**). Gene set enrichment analysis further identified enrichment of biological pathways associated with retinoblastoma transcriptome (**Figure S1C**). The heatmap (**Figure 5B**) revealed two major expression patterns. Genes associated with normal retinal identity and photoreceptor differentiation, including GNAT1, VSX2, PDE6B, RHO, GUCA1A, NRL, CARBP-1 and RBP3 showed lower expression in retinoblastoma tumours as compared to control retinal samples. Unsupervised hierarchical clustering of selected differentially expressed clearly separated control retinal tissues from retinoblastoma samples (**Figure 5C**). These patterns suggest that RB tumours are associated with loss or reduction of normal retinal differentiation associated transcriptional programs (27)

**Figure 5.**
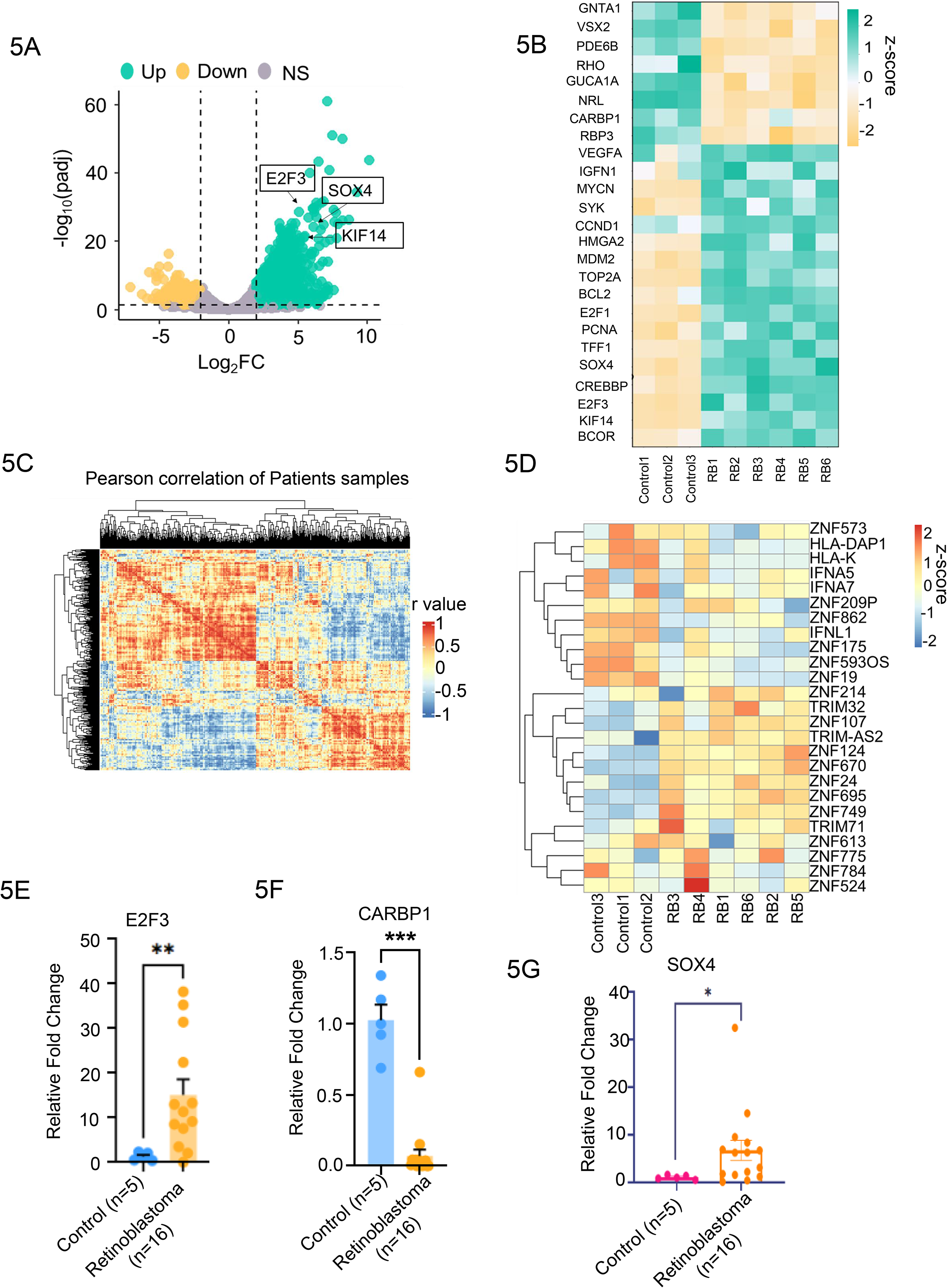
Retinoblastoma exhibits widespread transcriptional reprogramming associated with LINE-1 activation. **A.** The volcano plot highlights the distribution of significantly upregulated, downregulated, and non-significant (NS) genes. A total of 3,622 genes are upregulated, 1,994 are downregulated across the 6 patient samples compared to 3 controls. The x-axis represents the log₂ fold change (log₂FC), and the y-axis represents the -log₁₀ adjusted P value, reflecting both the magnitude and significance of differential expression. **B.** The heatmap visualizes the expression patterns of the top 25 genes with the most significant differential expression across 3 control and 6 patient samples. Upregulated genes are shown in green, while downregulated genes appear in yellow. Genes with no significant change in expression are represented in white. Each row corresponds to a single gene, and each column represents an individual sample. The color codes shows the level of gene expression ( z-score). **C.** Pearson correlation analysis was performed using all differentially expressed genes (DEGs) from six retinoblastoma (RB) patient samples. Significant positively correlated clusters are highlighted in red, while significant negatively correlated clusters are shown in light blue. **D.** The heatmap displays differential expression levels of the top 25 TEs repressor genes between 3 control and 6 RB patient’s sample. The x-axis shows the sample group and the y-axis shows the individuals genes. The colour codes ranges from red to blue. The red colour represents up regulated, blue colour represents down regulated genes and intermediate shades represents no change. **E-G.** mRNA quantification was performed via qRT-PCR using LINE1-specific primers that detect consensus sequences of E2F3, and CRABP1, SOX4, in control (n=5) and Retinoblastoma (n=16) tissue samples. Transcript levels were normalized to GAPDH, and relative fold changes were calculated relative to the control. Individual dots indicate biological replicates . Error bars show the standard error of the mean. *p<0.05, **p<0.01, ***p<0.01, statistical significance was determined using a two-tailed Mann-Whitney U test.

In contrast, several genes associated with proliferation, cell-cycle progression, chromatin regulation, survival, and oncogenic signalling showed increased expression in retinoblastoma samples compared with controls. These included MYCN, CCND1, HMGA2, MDM2, TOP2A, BCL2, E2F1, PCNA, SOX4, CREBBP, E2F3, KIF14 and BCOR. The coordinated increase of these genes in RB samples is consistent with an altered tumour-associated transcriptional state involving proliferative and regulatory pathways (28).

We next investigated the expression of genes that are associated with regulation of transposable element. Heatmap analysis revealed a variable expression of various host regulatory factors across retinoblastoma patient samples, including genes linked with DNA methylation dynamics, RNA/DNA processing, and intrinsic restriction of retrotransposons (**Figure 5D, Figure S2A and B**). Several genes, including TET1, TET2, TREX1, RNASEH2A, RNASEH2B, SAMHD1, and ZC3HAV1, exhibited elevated expression in subsets of RB tumors, whereas members of the APOBEC3 family, MOV10, and TEX19 exhibited variation in the transcript levels across RB tumors.

These results indicate that LINE-1 derepression in retinoblastoma occurs within a heterogeneous host regulatory environment rather than through a uniform change in all retrotransposon silencing programs. Together with the increased expression of selected L1HS loci and altered expression of tumour associated genes, our results demonstrate that LINE-1 derepression is accompanied by wider transcriptional changes involving both host silencing pathways and oncogenic programs. We also performed qRT-PCR experiments to validate transcriptomic changes of E2F3, CARBP1, and SOX4 in an independent cohort of control retinal tissues (n=5) and retinoblastoma tumours (n=16). Quantitative RT-PCR confirmed differential expression of these genes, consistent with the RNA sequencing results (**Figure 5E-G**).

These results suggests that endogenous LINE-1 activation in retinoblastoma is accompanied by coordinated transcriptional reprogramming characterized by elevated proliferative signalling, loss of retinal differentiation, and alteration in the levels of multiple host pathways responsible for maintaining silencing of transposons.

### Result 5. LINE-1 loci exhibit reduced repressive chromatin signatures in retinoblastoma

To examine the epigenetic landscape associated with activated LINE-1 loci, we analyzed publicly available ChIP-seq datasets for the repressive histone marks H3K9me3 and H3K27me3 in IMR-90 cells (29) using the Integrative Genomics Viewer (IGV) (**Figure 6A**). A representative genomic region encompassing an activated L1HS locus at chr6:85,953,220–86,051,335 was visualized. This locus was expressed across all six RB patient samples in our RNA-seq analysis (**Figure 4B**). Across this representative locus, we observed reduced enrichment of the heterochromatin associated histone marks H3K9me3 and H3K27me3 compared with control regions, while the input signals remained comparable between samples. These observations are consistent with reduced repressive chromatin at a representative LINE-1 locus and support the epigenetic derepression of LINE-1 elements observed in retinoblastoma.

**Figure 6.**
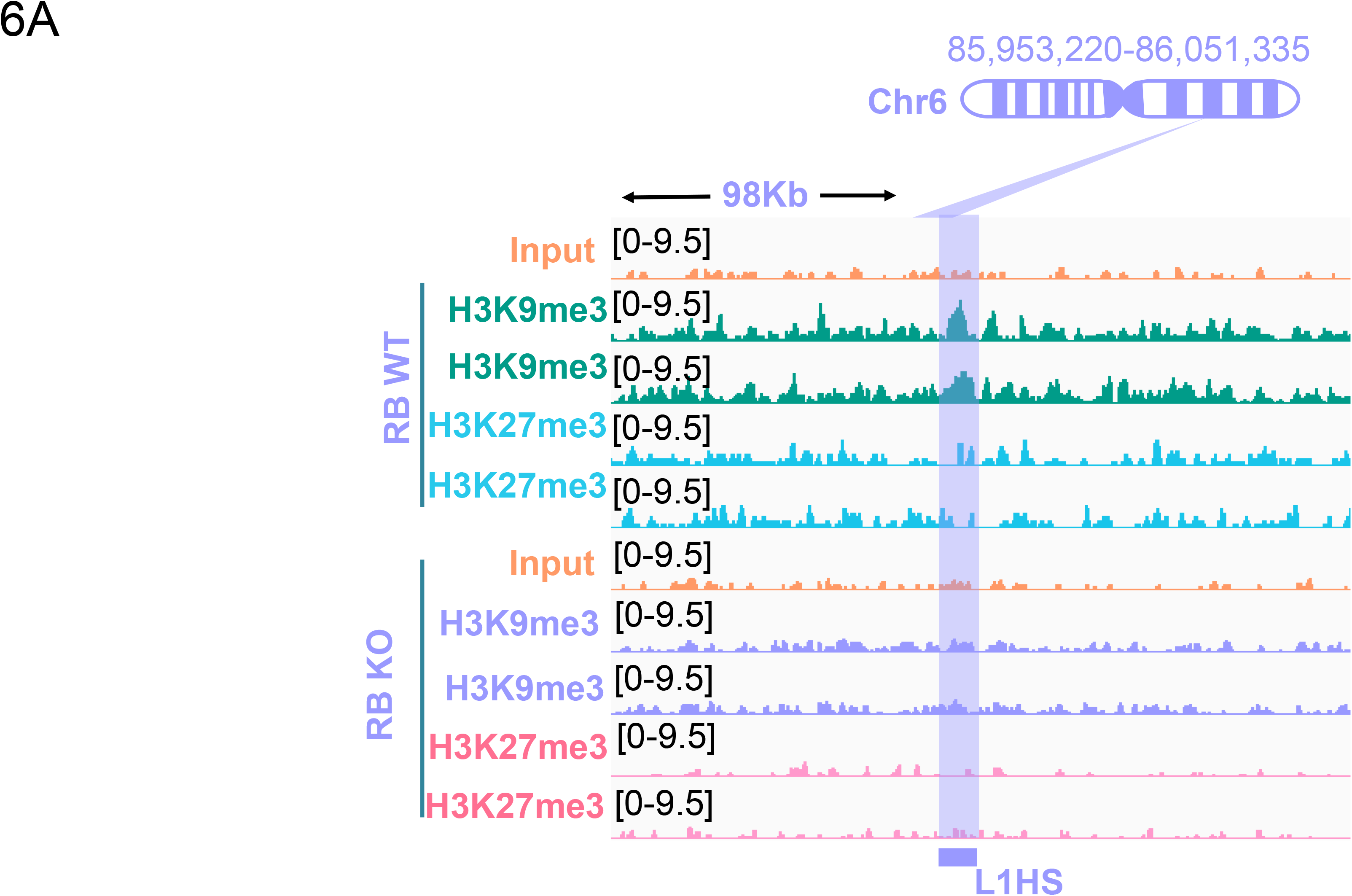
Loss of repressive chromatin marks accompanies activation of oncogenic transcriptional regulators in retinoblastoma. **A.** Interactive genome browser IGV tracks shows the chromatin landscape surrounding at L1HS on chromosome 6 (98 kb window). Input control and ChIP-seq profiles for repressive marks (H3K9me3 and H3K27me3) are shown.

## Discussion

Retinoblastoma is classically understood as a tumor initiated by loss of RB1 PMID: 27488068 (30), yet the clinical and molecular diversity of the disease indicates that additional layers of regulation may influence tumor behavior. In this study, we examined one such layer: the expression profile of transposable elements, particularly focussing on LINE-1 retrotransposons. LINE1 transposons is the autonomously active transposons in humans and are known to be reactivated in several human cancers (31,32). Our results suggest that LINE-1 derepression is part of the epigenetic and transcriptional landscape of retinoblastoma. We propose a model in which relaxation of repetitive element repression accompanies broader chromatin and gene expression changes in RB. However, further investigation is required to determine whether LINE-1 acts as a driver or promoter of these genomic and epigenomic changes in Retinoblastoma.

A key insight of our study is that Retinoblastoma mediated transposon silencing may constitute an important tumor suppressive axis in addition to well-established role of RB1 regulation on cell-cycle. RB1 is known to contribute to chromatin organization and transcriptional repression (33),and its loss may causes a permissive environment in which normally silenced genomic regions become transcriptionally active. In this scenario, LINE-1 promoter hypomethylation and ORF1p detection validate that at least a few subset of LINE-1 elements escape repression in RB tissues. This is important because LINE-1 activation has been shown to be associated with genome stability and cellular stress responses (34). However, whether these LINE-1 associated changes are functionally involved in RB progression or represent a consequence of broader epigenetic deregulation remains to be investigated.

Our findings that evolutionarily young L1Hs are preferentially represented among the altered LINE-1 signals is particularly important. The activation of younger LINE-1 subfamilies, including L1HS and related elements, suggests that the transcriptional relaxation observed in RB may lead to LINE-1 copies with higher residual activity. However, our study do not demonstrate L1 retrotransposition activity or novel somatic insertions. Future studies using long-read sequencing may be required to determine whether the observed transcriptional activation leads to retrotransposition dependent consequences.

The transcriptomic changes observed in RB also indicate to a broader remodeling of tumor-associated gene expression programs. Many of these genes are involved in chromatin regulation, cell-cycle control and retinal developmental pathways. These genes are relevant with the biology of RB as a developmental tumor in which proliferation, differentiation and epigenetic regulation are tightly linked. Although LINE-1 activation and host gene dysregulation occur together in the tumor samples, the present study does not establish a direct regulatory relationship between these observations.

The integration of publicly available chromatin datasets provides additional context for interpreting the transcriptomic findings. We observed reduced enrichment of the repressive histone marks H3K9me3 and H3K27me3 at an L1HS locus that was consistently derepressed across all six RB patient samples. This suggests that LINE-1 activation in RB may be associated with relaxation of repressive chromatin at selected LINE-1 loci, in addition to increased transcript expression. However, because these chromatin datasets were not generated from the same patient samples analyzed in this study, these cannot be used as direct evidence of chromatin changes in our cohort.

The strength of our work is the use of multiple orthogonal approaches to examine LINE-1 deregulation in patient derived RB tumour samples. This study combines transcriptomic analysis with qRT-PCR validation, LINE-1 promoter methylation profiling, transposable element expression analysis and immunohistochemical staining of ORF1p. To our knowledge, this represents the first transcriptomic analysis of retinoblastoma patient samples from an Indian cohort. This is significant because retinoblastoma is a rare pediatric tumor, and access to well annotated patient specimen remains limited. Population specific studies are important because stage at diagnosis, treatment access, clinical presentation and outcome can vary across healthcare settings.

The study also has limitations that should be considered. The cohort size is modest, and different molecular assays were performed on partially overlapping sample sets depending on tissue availability and sample quality. A major limitation is the restricted availability of appropriate retinal control samples. Obtaining healthy retinal tissue from individuals without retinoblastoma is ethically and practically challenging, especially in pediatric settings. As a result, the number of control retinal samples available for comparison was limited. Another limitation is that the present study remains primarily observational. While the data identify LINE-1 hypomethylation, LINE-1 transcript expression and ORF1p accumulation as characteristics associated with RB tissues, they do not prove that LINE-1 activity drives tumor initiation, progression or chromatin remodeling. Functional experiments will be needed to address causality.

Overall, this study expands the molecular view of retinoblastoma by highlighting dysregulated LINE-1 as part of its epigenetic and transcriptional landscape. The findings support a model in which loss of repetitive element repression accompanies broader chromatin and gene expression changes in RB. While the clinical and functional consequences of this deregulation remain to be defined, the study provides a foundation for future work investigating whether LINE-1 associated mechanisms contribute to tumor heterogeneity, genome instability or disease severity in this rare pediatric ocular malignancy.

## Material and Methods

### Patient Recruitment

All RB patients undergoing enucleation at a tertiary eye care ocular oncology centre were included in the study. The inclusion criteria consisted of tumor tissue from RB patients post-enucleation, obtained with written consent from parents/guardians; detailed demographics are provided in the Supplementary Table. Institutional ethics approval was obtained and all procedures adhered to the Declaration of Helsinki and ARVO best-practice guidelines for human eye tissue research. Ethical approval was obtained from the institutional review board (IRB/2025/MAR/08). Exclusion criteria included patients who did not undergo enucleation as part of their treatment, patients with parents/guardians who did not consent to the study, and patients lost to follow-up after enucleation.

### Total RNA Isolation

RNA was isolated from retinoblastoma patient enucleated eyeballs and from normal retinas from enucleated eyeballs using a kit-based mansufacturer’s protocol (All prep DNA/RNA/Protein Mini Kit QIAGEN kit Cat No. 80004 / Qiagen RNA easy kit No. 74104).

### Real-Time PCR experiments (qPCR)

A Qiagen Reverse transcriptase two-step cDNA kit (Catalog no.205311, Qiagen) was used to synthesize cDNA from RNA. qPCR was performed using cDNA and PowerUp SYBR Green PCR master mix (Catalog no. A25742, Applied Biosystems by Thermo Fisher Scientific brand) on the Azure Cielo Real-time PCR system thermocycler (Catalog no.Q16-1347, Azure Cielo 6, Azure Biosystems, Inc). PCR parameters were 95 °C for 3 min, followed by 40 cycles of denaturation at 95 °C for 10 s, annealing at 60 °C for 20 s, and extension at 60 °C for 5 s, with a final step at 95 °C for 1 min. CT values were normalized with GAPDH housekeeping genes, and the relative fold change was calculated using retinal control tissues.

### Immunohistochemistry

Enucleated eyeballs for retinoblastoma cases were carefully excised and fixed in formalin. The specimens were processed through standard histopathological protocols, including dehydration, clearing, and paraffin embedding. Serial sections of 2.5 micrometers thickness were cut using a microtome and mounted on glass slides. The sections were then stained with Hematoxylin and Eosin (H&E) to facilitate detailed examination of tumor morphology, cellular architecture, and infiltration patterns. Slides with well-preserved tumor tissue, exhibiting clear morphological features, were selected for microscopic analysis to enhance understanding of tumor characteristics. Oral Squamous Cell Carcinoma was used as a positive control for LINE1 expression. Tissues were fixed in 10% formalin, embedded in paraffin, sectioned, and mounted on positively charged slides. Slides were baked at 55–60°C for 45 minutes, deparaffinized with xylene, and rehydrated through a graded ethanol series. Heat-induced epitope retrieval was performed using sodium citrate buffer preheated in a water bath at 98°C, followed by washing with 1X TBS (Tris-buffered saline). Endogenous peroxidase activity was quenched with the kit’s hydrogen peroxide for 15 min in a humid chamber, and the slides were then washed with 1X TBS. Slides were then blocked with 1% BSA in TBS for 30 mins at room temperature in a humid chamber. Slides were incubated overnight at 4°C with anti-LINE1 ORF1 antibody (Merck Millipore, MABC1152; dilution 1:500). Next, they were washed with 1X TBS and incubated with DAB chromogen from the PathnSitu kit (Catalog no. PEH002, PathnSitu Biotechnologies), followed by hematoxylin counterstaining and DPX mounting.

### Genomic DNA isolation

Retinoblastoma tumor tissue and retinal tissue were collected post-enucleation in 1.5ml MCT and washed with phosphate buffer saline (PBS), followed by homogenization of the sample in liquid nitrogen using a homogenizer. From homogenized tissue, genomic DNA was extracted using as per the manufacturer’s protocol (All prep DNA/RNA/Protein Mini Kit QIAGEN kit Cat No. 80004/ Blood &Tissue DNeasy No. 69504)

### Methylation analysis

LINE-1 promoter methylation was performed using Bisulfite conversion of gDNA (1000 ng) isolated from tissue specimens, following the manufacturer’s instructions for the EZ DNA Methylation™ Kit (Cat. No. D5005). A 363 bp sequence within the L1 5’ UTR region, containing twenty CpG dinucleotides, was amplified using a methylated primer set (Forward: 5’-AAGGGGTTAGGGAGTTTTTTT-3’ and Reverse: 5’-TATCTATACCCTACCCCCAAAA-3’), as per the sequence and base pair length reported in the paper and calibrated according to our laboratory protocols.

A 50 μl PCR reaction was prepared using GeneI master mix( Catlog no. - 060220003A) and 200 ng of bisulfite-treated genomic DNA. Untreated genomic DNA served as a negative control. The lack of PCR amplification in the untreated sample confirmed complete conversion of cytosines. PCR amplification conditions included an initial denaturation at 94 °C for 30 seconds, followed by 30 cycles at 94 °C for 20 seconds, 54 °C for 30 seconds, 72 °C for 60 seconds, and a final extension at 72 °C for 5 minutes. PCR products were separated on a 1.2% agarose gel, excised, and purified using a gel extraction kit (Thermo Scientific, Catalog No. K0701). The purified products were cloned into the pGEM-T vector (Promega, Catalog No. A3600), followed by transformation into competent cells and blue-white screening. Plasmid DNA was extracted from white colonies using a Qiagen miniprep kit. Insert verification was performed by electrophoresis in 1.5% agarose gel, and five clones per sample were sequenced using Sanger sequencing with T7 primer. Sequences were manually inspected to determine methylation status at CpG sites, noting whether cytosines remained intact (methylated) or were converted (unmethylated). Methylation analysis was conducted using QUMA and BioEdit software to quantify methylation levels.

### Bulk RNA Sequencing

Total RNA was quantified using a Thermo ScientificTM NanoDrop and outsourced for sequencing. The RNA-seq analysis encompasses quality control of raw and trimmed reads, alignment statistics, and differential gene expression analysis. All quality metrics indicate a successful run with good quality sequencing data and reliable downstream analysis results.

Wet Lab: Libraries were prepared using IAGEN’s IAseq FastSelect RNA Lib HMR Kit (24) as per the protocol for totalRNA library. Library C was performed using BioAnalyzer (Agilent) and was quantified using UBIT Fluorometer (LifeTech). Libraries have been normalized and pooled for sequencing on the NovaSeq 6000 (Illumina) in accordance with sequencing guidelines.

### RNASeq Analysis (in house)

Raw sequencing data quality was initially assessed using FastQC (v0.12.1) (fastqc <input_file> -o <output_directory>) (35). Only high-quality reads were aligned to the human reference genome (hg38) using Bowtie2 (v2.5.4) (bowtie2 -x <reference_genome_index> -1 <R1> -2 <R2> -S <output.sam> --threads <N>) (36). The SAM files were converted to BAM format, sorted, and indexed using SAMtools (v1.19.2) (samtools view -bS <input.sam> -o <output.bam> and samtools sort <input.bam> -o <output.sorted.bam>) (37). Quantification of both genes and transposable elements (TEs) was performed using TEcount from the TEtranscripts package (v2.2.4) (TEcount -b <file> --format BAM --mode multi --stranded no --GTF <gene_annotation.gtf> --TE <TE_annotation.gtf> --project <sample_name> --sortByPos) (38). TEcount generated count tables containing read counts for both genes and TEs for each sample. Gene and TE count matrices were subsequently separated using inhouse custom Python scripts for downstream analyses. Differential expression analysis was performed using DESeq2 (v4.3.3) in R (v4.5.0) (39). Genes and transposable elements with an absolute log2 fold change greater than 1 and an adjusted p-value (Benjamini-Hochberg corrected) less than 0.05 were considered significantly differentially expressed between control and retinoblastoma (RB) patient samples. For downstream visualization and clustering analyses, count data were transformed using the variance stabilizing transformation (VST) implemented in DESeq2, followed by z-score normalization where applicable. All statistical plots and visualizations were generated using ggplot2 (v4.3.1) in R (40). For the locus specific LINE1 analysis we used TElocal tools provided by TEtranscript. The gene set enrichment analysis (GSEA) was done using ShinyGO (v0.85)(41).

### ChIPseq Analysis

The raw data were downloaded from the NCBI Gene Expression Omnibus (GEO) database and retrieved using the SRA Toolkit (v3.4.1)(42) . SRA files were converted into paired-end FASTQ files using the option (fastq-dump --split-files <input.sra>). Data preprocessing was performed following the same pipeline described above for RNA-seq analysis. For ChIP-seq analysis, reads were aligned to the human reference genome (hg38) using Bowtie2. The SAM files were converted to BAM format and processed using SAMtools (v). BigWig files were generated using deepTools bamCoverage (v3.5.5) with FPKM normalization (bamCoverage -b <input.bam> -o <output.bw> --normalizeUsing FPKM -p <threads>)(43). Peak calling was performed using MACS3 (v3.0.2) (macs3 callpeak -t <treatment.bam> -c <control.bam> -f BAM -g hs -n <output_file>), generating narrowPeak files for downstream analyses (44). Signal enrichment profiles and heatmaps were generated using deepTools. The computeMatrix function was used to calculate signal matrices, and plotHeatmap was used to visualize the distribution of ChIP-seq signals across genomic regions of interest. BigWig tracks for both H3K27me3 and H3K9me3 were visualized using the Integrative Genomics Viewer (IGV) (v2.16.0) (45).

### Statistics

Statistical analyses used Prism 7.01 (GraphPad Inc.), with Student’s two tailed t test for two datasets and analysis of variance for multiple comparisons; p < 0.05 was considered significant.

## Data Availability

ChIP-Seq datasets described in our study can be found in NCBI GEO id (GSE173125).

## Funding and Additional Information

This work was supported by the DBT/Wellcome Trust India Alliance fellowship (IA/I/22/2/506501) to Bhavana Tiwari at IISER Berhmpur This work was also partly supported by the ANRF CRG/2023/007/7033 to Bhavana Tiwari at IISER Berhampur. This work is supported by Indian Council of Medical Research grant (EMDR/IG/11/2024-01907 & 2021-15609F1) to Dr Virender Singh Sangwan and Dr Anil Tiwari at Dr Shroff’s Charity Eye Hospital, Delhi

## CRedit authorship contribution statement

**Aman Verma:** Investigation, Data curation, Formal analysis, methodology, writing original draft

**Ria Sachdeva:** Clinical Data curation, patient recruitment

**Akhil Varshney:** Conceptualization, Investigation.

**Virender Singh Sangwan:** Resources, Patient recruitment, Clinical supervision.

**Sima Das:** Patient recruitment, Sample collection, Clinical data curation, Investigation.

**Sugam Kumar Patel:** Software and Formal analysis, Data visualization, Writing - original draft.

**Pratyashaa Paul:** Methodology, Visualization.

**Bhavana Tiwari:** Conceptualization, Supervision, Writing - original draft, Writing – review & editing, Funding acquisition.

**Anil Tiwari:** Supervision, Project administration, Funding acquisition, Writing - review & editing.

## Acknowledgements

We thank the Central Instrumentation Facility and the Department of Biological Sciences at IISER Berhampur for providing infrastructure support. We also acknowledge the Eicher Shroff Centre for Stem Cell Research for institutional support and Dr Chhavi Gupta, Dr Anureet Kaur for the clinical support. Finally, we sincerely thank the patients and their families for participating in this study. This work forms part of the Ph.D. thesis of Aman Verma (registered with Manipal Academy of Higher Education, Enrollment No. 239000027)

## References

1. Dimaras H, Corson TW, Cobrinik D, White A, Zhao J, Munier FL, et al. Retinoblastoma. Nat Rev Dis Primer. 2015 Aug 27;1(1):15021. doi:10.1038/nrdp.2015.21

2. McEvoy JD, Dyer MA. Genetic and Epigenetic Discoveries in Human Retinoblastoma. Crit Rev Oncog. 2015;20(3–4):217–25. doi:10.1615/critrevoncog.2015013711 PubMed PMID: 26349417; PubMed Central PMCID: PMC5458782.

3. Nag A, Khetan V. Retinoblastoma - A comprehensive review, update and recent advances. Indian J Ophthalmol. 2024 Jun 1;72(6):778–88. doi:10.4103/IJO.IJO_2414_23 PubMed PMID: 38804799; PubMed Central PMCID: PMC11232864.

4. Shields CL, Bas Z, Laiton A, Silva AMV, Sheikh A, Lally SE, et al. Retinoblastoma: emerging concepts in genetics, global disease burden, chemotherapy outcomes, and psychological impact. Eye. 2023 Apr;37(5):815–22. doi:10.1038/s41433-022-01980-0 PubMed PMID: 35217824; PubMed Central PMCID: PMC8873344.

5. 5. Global Retinoblastoma Study Group. The Global Retinoblastoma Outcome Study: a prospective, cluster-based analysis of 4064 patients from 149 countries. Lancet Glob Health. 2022 Aug;10(8):e1128–40. doi:10.1016/S2214-109X(22)00250-9 PubMed PMID: 35839812; PubMed Central PMCID: PMC9397647.

6. Ratna R, Varshney A, Tibrewal S, Verma A, Majumdar A, Das S. Mapping RB1 gene mutations in retinoblastoma: a study of 200 cases from North India. Ophthalmic Genet. 2025 Dec;46(6):552–8. doi:10.1080/13816810.2025.2518136 PubMed PMID: 40534329.

7. Das S, Meel R, Mahajan A, Bansal R, Reddy VA, Prasad S, et al. Lag time for diagnosis and treatment in 1120 retinoblastoma children: Analysis from InPOG-RB-19-01. Indian J Ophthalmol. 2025 Aug 1;73(8):1124–31. doi:10.4103/IJO.IJO_3031_24 PubMed PMID: 40719713; PubMed Central PMCID: PMC12416608.

8. Dick FA, Rubin SM. Molecular mechanisms underlying RB protein function. Nat Rev Mol Cell Biol. 2013 May;14(5):297–306. doi:10.1038/nrm3567

9. Lee H, Gkotinakou IM, McGrath CG, Krishnan B, Animesh S, Salinas I, et al. RB loss modulates chromatin organization by regulating cohesin-dependent loops and enhancer-promoter interactions. Nat Commun. 2026 Apr 8;17(1):3696. doi:10.1038/s41467-026-71655-x PubMed PMID: 41951674; PubMed Central PMCID: PMC13103356.

10. Ishak CA, Marshall AE, Passos DT, White CR, Kim SJ, Cecchini MJ, et al. An RB-EZH2 Complex Mediates Silencing of Repetitive DNA Sequences. Mol Cell. 2016 Dec 15;64(6):1074–87. doi:10.1016/j.molcel.2016.10.021 PubMed PMID: 27889452; PubMed Central PMCID: PMC5340194.

11. Kazazian HH, Goodier JL. LINE Drive: Retrotransposition and Genome Instability. Cell. 2002 Aug 9;110(3):277–80. doi:10.1016/S0092-8674(02)00868-1

12. Brouha B, Schustak J, Badge RM, Lutz-Prigge S, Farley AH, Moran JV, et al. Hot L1s account for the bulk of retrotransposition in the human population. Proc Natl Acad Sci U S A. 2003 Apr 29;100(9):5280–5. doi:10.1073/pnas.0831042100 PubMed PMID: 12682288; PubMed Central PMCID: PMC154336.

13. Ardeljan D, Taylor MS, Ting DT, Burns KH. The human LINE-1 retrotransposon: an emerging biomarker of neoplasia. Clin Chem. 2017 Apr;63(4):816–22. doi:10.1373/clinchem.2016.257444 PubMed PMID: 28188229; PubMed Central PMCID: PMC6177209.

14. Rodić N, Burns KH. Long interspersed element-1 (LINE-1): passenger or driver in human neoplasms? PLoS Genet. 2013 Mar;9(3):e1003402. doi:10.1371/journal.pgen.1003402 PubMed PMID: 23555307; PubMed Central PMCID: PMC3610623.

15. Rodriguez-Martin B, Alvarez EG, Baez-Ortega A, Zamora J, Supek F, Demeulemeester J, et al. Pan-cancer analysis of whole genomes identifies driver rearrangements promoted by LINE-1 retrotransposition. Nat Genet. 2020 Mar;52(3):306–19. doi:10.1038/s41588-019-0562-0

16. Tiwari B, Jones AE, Caillet CJ, Das S, Royer SK, Abrams JM. p53 directly represses human LINE1 transposons. Genes Dev. 2020 Nov 1;34(21–22):1439–51. doi:10.1101/gad.343186.120 PubMed PMID: 33060137; PubMed Central PMCID: PMC7608743.

17. Paul P, Kumar A, Parida AS, De AK, Bhadke G, Khatua S, et al. p53-mediated regulation of LINE1 retrotransposon-derived R-loops. J Biol Chem. 2025 Mar;301(3):108200. doi:10.1016/j.jbc.2025.108200 PubMed PMID: 39828096; PubMed Central PMCID: PMC11903798.

18. Tiwari B, Jones AE, Abrams JM. Transposons, p53 and Genome Security. Trends Genet TIG. 2018 Nov;34(11):846–55. doi:10.1016/j.tig.2018.08.003 PubMed PMID: 30195581; PubMed Central PMCID: PMC6260979.

19. Beck CR, Garcia-Perez JL, Badge RM, Moran JV. LINE-1 elements in structural variation and disease. Annu Rev Genomics Hum Genet. 2011;12:187–215. doi:10.1146/annurev-genom-082509-141802 PubMed PMID: 21801021; PubMed Central PMCID: PMC4124830.

20. McClendon AK, Dean JL, Ertel A, Fu Z, Rivadeneira DB, Reed CA, et al. RB and p53 cooperate to prevent liver tumorigenesis in response to tissue damage. Gastroenterology. 2011 Oct;141(4):1439–50. doi:10.1053/j.gastro.2011.06.046 PubMed PMID: 21704587; PubMed Central PMCID: PMC4562673.

21. Knudsen ES, Nambiar R, Rosario SR, Smiraglia DJ, Goodrich DW, Witkiewicz AK. Pan-cancer molecular analysis of the RB tumor suppressor pathway. Commun Biol. 2020 Apr 2;3:158. doi:10.1038/s42003-020-0873-9 PubMed PMID: 32242058; PubMed Central PMCID: PMC7118159.

22. Vandel L, Nicolas E, Vaute O, Ferreira R, Ait-Si-Ali S, Trouche D. Transcriptional repression by the retinoblastoma protein through the recruitment of a histone methyltransferase. Mol Cell Biol. 2001 Oct;21(19):6484–94. doi:10.1128/MCB.21.19.6484-6494.2001 PubMed PMID: 11533237; PubMed Central PMCID: PMC99795.

23. Brehm A, Miska EA, McCance DJ, Reid JL, Bannister AJ, Kouzarides T. Retinoblastoma protein recruits histone deacetylase to repress transcription. Nature. 1998 Feb 5;391(6667):597–601. doi:10.1038/35404 PubMed PMID: 9468139.

24. Siddiqui H, Fox SR, Gunawardena RW, Knudsen ES. Loss of RB compromises specific heterochromatin modifications and modulates HP1alpha dynamics. J Cell Physiol. 2007 Apr;211(1):131–7. doi:10.1002/jcp.20913 PubMed PMID: 17245754.

25. Montoya-Durango DE, Liu Y, Teneng I, Kalbfleisch T, Lacy ME, Steffen MC, et al. Epigenetic control of mammalian LINE-1 retrotransposon by retinoblastoma proteins. Mutat Res - Fundam Mol Mech Mutagen. 2009 Jun 1;665(1):20–8. doi:10.1016/j.mrfmmm.2009.02.011

26. You JS, Pierce S, Liang G, Jones PA. Roles of transposable elements and DNA methylation in the formation of CpG islands and CpG-depleted regulatory elements. Proc Natl Acad Sci U S A. 122(43):e2502963122. doi:10.1073/pnas.2502963122 PubMed PMID: 41134632; PubMed Central PMCID: PMC12582260.

27. Burkhart DL, Sage J. Cellular mechanisms of tumour suppression by the retinoblastoma gene. Nat Rev Cancer. 2008 Sep;8(9):671–82. doi:10.1038/nrc2399 PubMed PMID: 18650841; PubMed Central PMCID: PMC6996492.

28. Gómez-Romero L, Alvarez-Suarez DE, Hernández-Lemus E, Ponce-Castañeda MV, Tovar H. The regulatory landscape of retinoblastoma: a pathway analysis perspective. R Soc Open Sci. 9(5):220031. doi:10.1098/rsos.220031 PubMed PMID: 35620002; PubMed Central PMCID: PMC9114937.

29. Wong KM, King DA, Schwartz EK, Herrera RE, Morrison AJ. Retinoblastoma protein regulates carcinogen susceptibility at heterochromatic cancer driver loci. Life Sci Alliance. 2022 Apr 1;5(4). doi:10.26508/lsa.202101134 PubMed PMID: 34983823.

30. Mallipatna A, Marino M, Singh AD. Genetics of Retinoblastoma. Asia-Pac J Ophthalmol. 2016 Jul 1;5(4):260–4. doi:10.1097/APO.0000000000000219

31. Mendez-Dorantes C, Burns KH. LINE-1 retrotransposition and its deregulation in cancers: implications for therapeutic opportunities. Genes Dev. 2023;37(21–24):948– 67. doi:10.1101/gad.351051.123 PubMed PMID: 38092519; PubMed Central PMCID: PMC10760644.

32. Burns KH. Transposable elements in cancer. Nat Rev Cancer. 2017 Jul;17(7):415–24. doi:10.1038/nrc.2017.35

33. Chinnam M, Goodrich DW. RB1, development, and cancer. Curr Top Dev Biol. 2011;94:129–69. doi:10.1016/B978-0-12-380916-2.00005-X PubMed PMID: 21295686; PubMed Central PMCID: PMC3691055.

34. Zhang X, Zhang R, Yu J. New Understanding of the Relevant Role of LINE-1 Retrotransposition in Human Disease and Immune Modulation. Front Cell Dev Biol. 2020;8:657. doi:10.3389/fcell.2020.00657 PubMed PMID: 32850797; PubMed Central PMCID: PMC7426637.

35. 35. Andrews S. FastQC: A Quality Control Tool for High Throughput Sequence Data [Internet]. Babraham Bioinformatics; 2010. Available from: https://www.bioinformatics.babraham.ac.uk/projects/fastqc/

36. Langmead B, Salzberg SL. Fast gapped-read alignment with Bowtie 2. Nat Methods. 2012 Mar 4;9(4):357–9. doi:10.1038/nmeth.1923 PubMed PMID: 22388286; PubMed Central PMCID: PMC3322381.

37. Li H, Handsaker B, Wysoker A, Fennell T, Ruan J, Homer N, et al. The Sequence Alignment/Map format and SAMtools. Bioinformatics. 2009 Aug 15;25(16):2078–9. doi:10.1093/bioinformatics/btp352 PubMed PMID: 19505943; PubMed Central PMCID: PMC2723002.

38. 38. Jin Y, Tam OH, Paniagua E, Hammell M. TEtranscripts: a package for including transposable elements in differential expression analysis of RNA-seq datasets. Bioinformatics. 2015 Nov 15;31(22):3593–9. doi:10.1093/bioinformatics/btv422 PubMed PMID: 26206304; PubMed Central PMCID: PMC4757950.

39. Love MI, Huber W, Anders S. Moderated estimation of fold change and dispersion for RNA-seq data with DESeq2. Genome Biol. 2014;15(12):550. doi:10.1186/s13059-014-0550-8 PubMed PMID: 25516281; PubMed Central PMCID: PMC4302049.

40. Wickham H. ggplot2 [Internet]. doi:10.1002/wics.147

41. Ge SX, Jung D, Yao R. ShinyGO: a graphical gene-set enrichment tool for animals and plants. Bioinformatics. 2020 Apr 1;36(8):2628–9. doi:10.1093/bioinformatics/btz931

42. 42. Maurya A, Szymanski M, Karlowski WM. ARA: a flexible pipeline for automated exploration of NCBI SRA datasets. GigaScience. 2023 Jan 1;12:giad067. doi:10.1093/gigascience/giad067

43. Ramírez F, Dündar F, Diehl S, Grüning BA, Manke T. deepTools: a flexible platform for exploring deep-sequencing data. Nucleic Acids Res. 2014 Jul 1;42(W1):W187–91. doi:10.1093/nar/gku365

44. Zhang Y, Liu T, Meyer CA, Eeckhoute J, Johnson DS, Bernstein BE, et al. Model-based Analysis of ChIP-Seq (MACS). Genome Biol. 2008;9(9):R137. doi:10.1186/gb-2008-9-9-r137 PubMed PMID: 18798982; PubMed Central PMCID: PMC2592715.

45. Robinson JT, Thorvaldsdóttir H, Winckler W, Guttman M, Lander ES, Getz G, et al. Integrative Genomics Viewer. Nat Biotechnol. 2011 Jan;29(1):24–6. doi:10.1038/nbt.1754 PubMed PMID: 21221095; PubMed Central PMCID: PMC3346182.

