## Supplemental files for "LINE-1 transposon derepression and epigenetic remodeling in Retinoblastoma"

Supplementary Figure 1

S1A

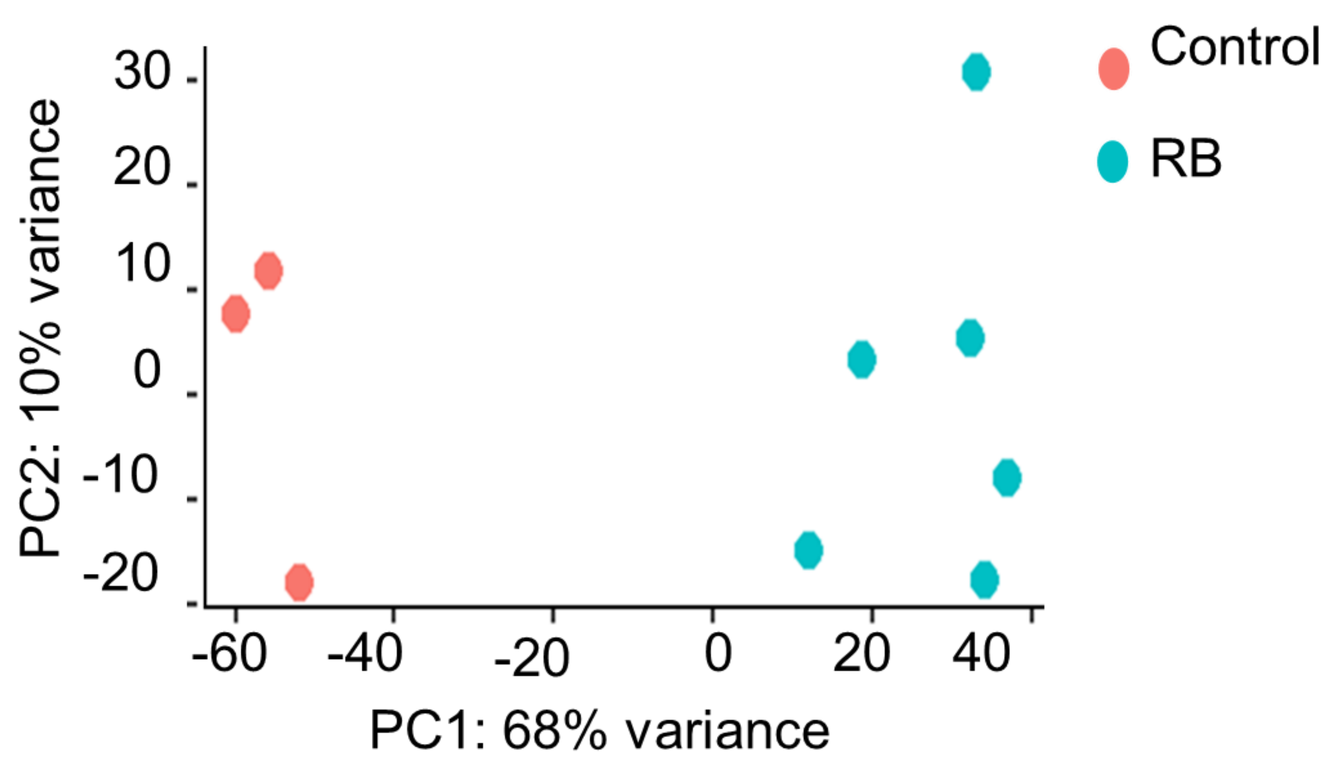


S1B

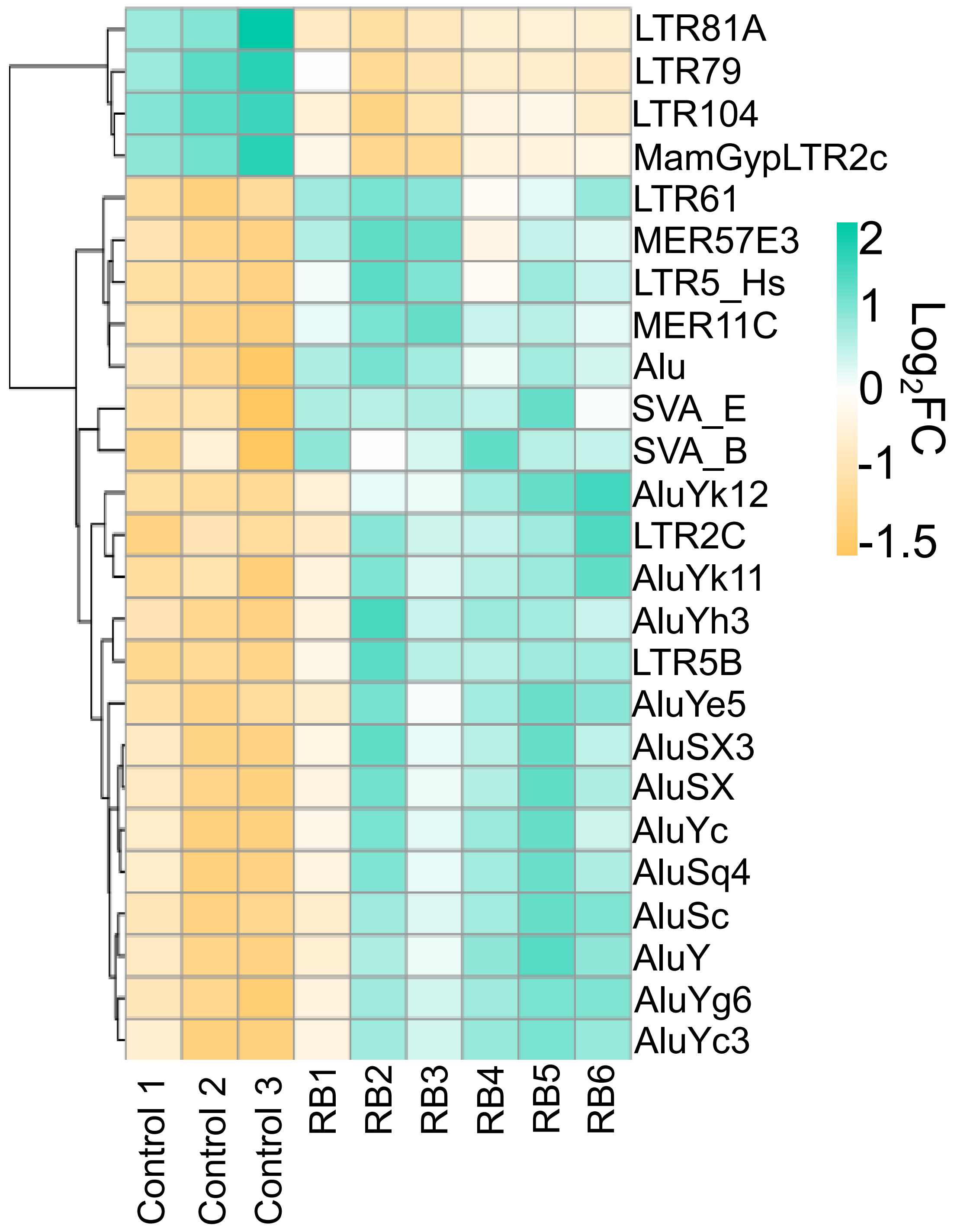


S1C

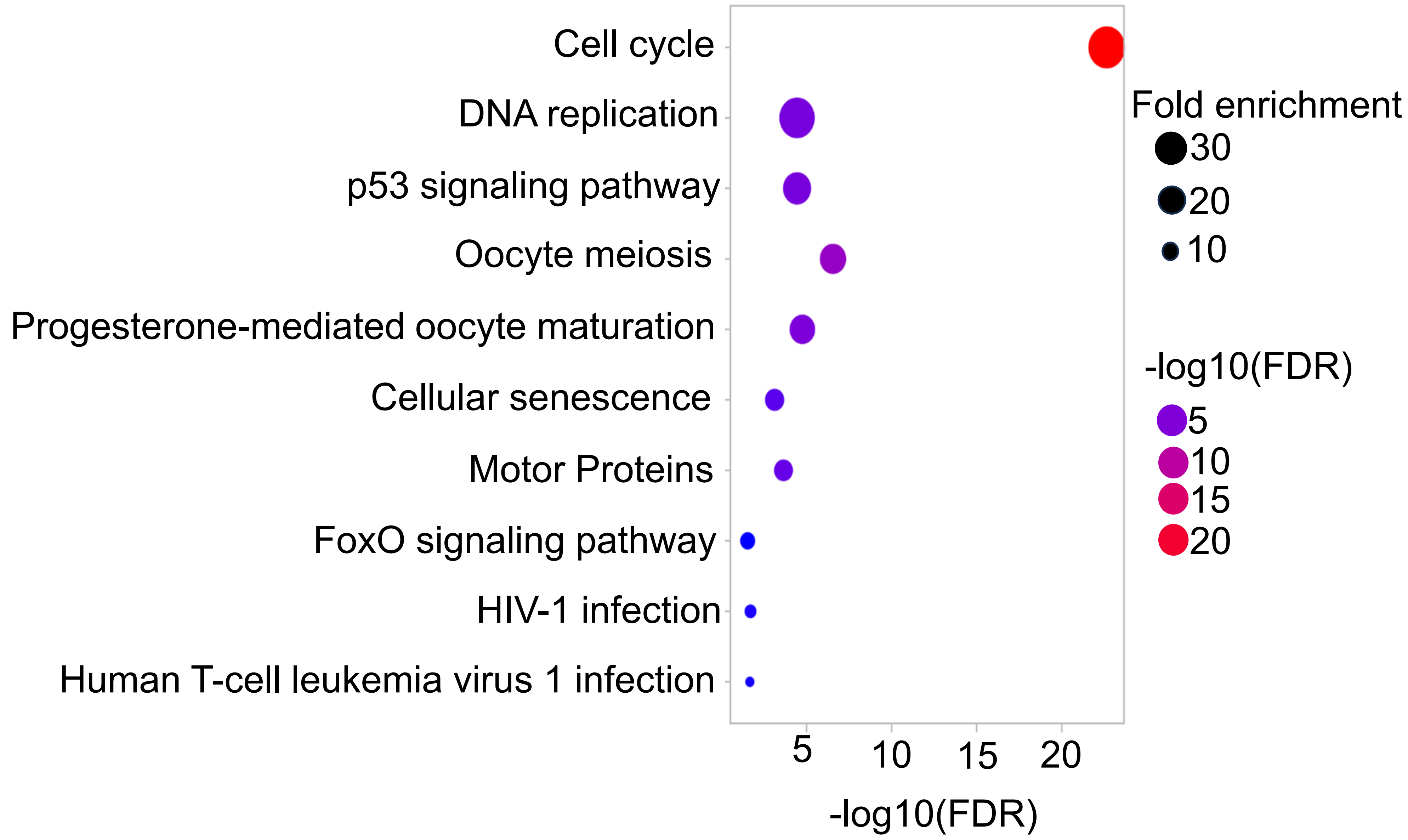

**Figure S1A.** The PCA plot shows distinction between the 3 control samples (red dots) and the 6 patient samples (green dots). Principal Component 1 (PC1) explains 68% of the total variance, while Principal Component 2 (PC2) accounts for an additional 10%.

**Figure S1B.**The heatmap represents top 25 differentially expressed TEs family except LINE1 in control and RB patient's sample. The green colour shows up regulated, yellow colour shows the down regulated and the intermediate shades show no change. The x-axis represents samples group and the y-axis represents individuals TEs family.

**Figure S1C.** The bubble plots represents Gene Set Enrichment Analysis (GSEA) of top enrichment genes. The x-axis shows the $-\log_{10}$(FDR) on the y-axis shows particular pathway name, the bubble size indicates fold enrichment.

Supplementary Figure 2

S2A

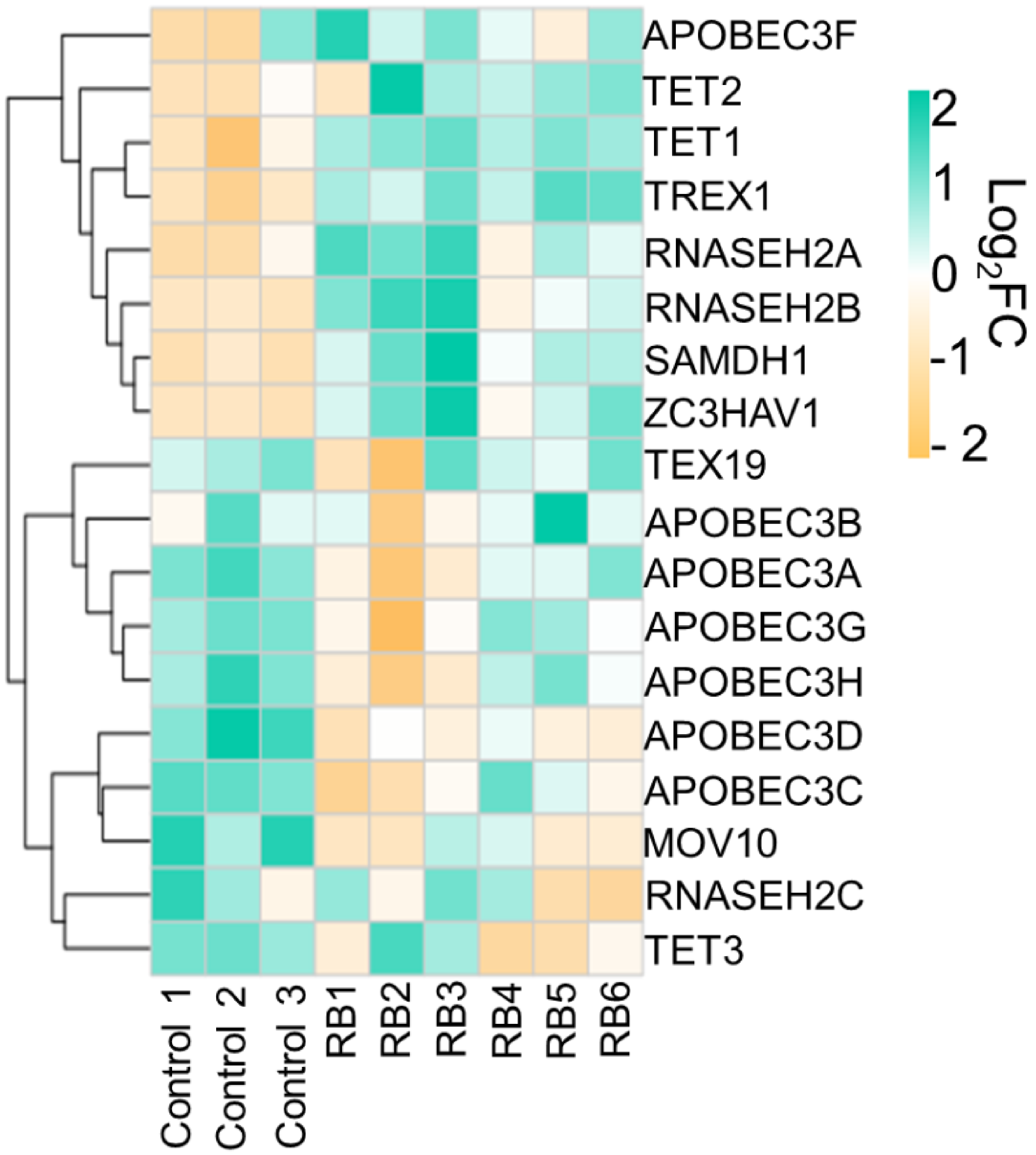


S2B

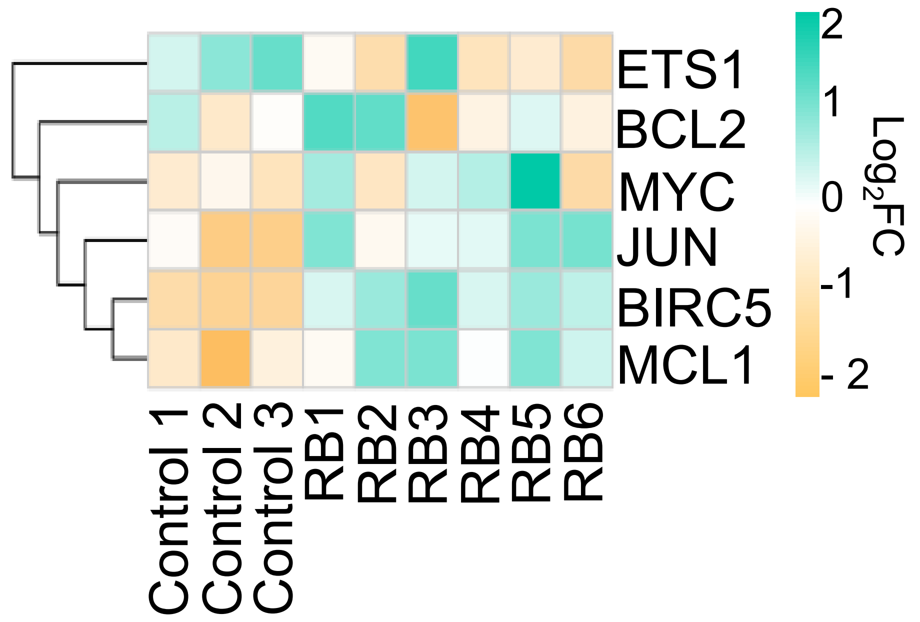

**Figure S2A** The heatmap represents transcription factor genes associated with RB patient's sample. The x-axis shows the sample group and y-axis show individuals genes. The color codes ranges from green (up regulated) yellow (down regulated) and intermediate shades shows no change.

**Figure S2B** The heatmap represents restriction factor genes associated with RB patient's sample. The x-axis shows the sample group and y-axis show individuals genes. The color codes ranges from green (up regulated) yellow (down regulated) and intermediate shades shows no change.
